# Nucleic acid quantification with amplicon yield in recombinase polymerase amplification

**DOI:** 10.1101/2022.06.28.497931

**Authors:** Priyanka Valloly, Rahul Roy

## Abstract

Amplification-based qPCR provides accurate and sensitive nucleic acid quantification. However, the requirement of temperature cycling and real-time monitoring limits its translation to different settings. Here, we adapted isothermal Recombinase Polymerase Amplification (RPA) reaction to develop a semi-quantitative method that relies on final amplicon yield to estimate initial target nucleic acid copy number. To achieve this, we developed a phenomenological model that captures the essential RPA dynamics. We identified reaction conditions that constrained the reaction yield corresponding to the starting DNA template concentration. We validated these predictions experimentally and show that the amplicon yields at the end of the RPA reaction correlates well to the starting DNA concentration while reducing non-specific amplification robustly. We demonstrate this approach termed here as quantitative endpoint RPA (qeRPA) to detect DNA over five log orders with detection limit of 100 molecules. Using a linear regression model with normalized endpoint intensity (NEI) standard curve, we estimate viral load from the serum of dengue-infected patients with comparable performance to qPCR. Hence, qeRPA can be employed for robust and sensitive nucleic acid estimation at close to room temperature without real-time monitoring and can be beneficial for field-deployment in limited-resource settings.

## Introduction

Nucleic acid quantification (NAQ) is crucial for gene expression analysis, detection of rare or dysfunctional cells, monitoring pathogens during infections, and assessing treatment outcomes.^1–3^ Quantitative Polymerase Chain Reaction (qPCR) has long been the gold standard for nucleic acid analysis and quantification.^4,5^ Since the key observable, threshold cycle (Ct) estimate is dependent on the efficiency and kinetics of the polymerization reaction, qPCR requires careful calibration and continuous monitoring of the amplification reaction. More importantly, the necessity for thermal cycling makes its implementation difficult in resource-limited settings. The recent advent of several isothermal nucleic acid amplification techniques (INAATs) alleviate the need for thermal cycling and hence offer promising alternatives to qPCR.^6^ INAATs like Loop Mediated Isothermal Amplification (LAMP),^7^ Helicase dependent amplification (HDA),^8^ Strand displacement amplification (SDA)^9^ can generate detectable amplicons in less than an hour. Using the time to reach amplicon levels beyond the threshold of detection, *τ_d_,* INAATs can also report estimates of initial DNA concentration.^10^ For example, LAMP based DNA detection with a dynamic range of 10 - 10^7^ copies/*μ*l has been reported earlier.^11^ However, rapid amplification (that limits tight temporal control), non specific products (that mandates use of specific probes) and reliance on real-time monitoring pose challenges for developing a robust and field-deployable DNA quantification method with INAATs.

Another promising INAAT, Recombinase Polymerase Amplification (RPA), works close to ambient temperature (37 - 42 °C) and requires two end-specific primers with a cocktail of recombinase, single-strand binding protein, and a strand displacement DNA polymerase.^12,13^ RPA reaction is sensitive yet rapid and can amplify even a single molecule of target DNA within 30 minutes to detectable levels.^14^ Yet, most successfully developed RPA assays are qualitative with positive/negative readout for target DNA and *τ_d_* based quantification from real-time RPA kinetic measurements has found limited success.^14–16^ This partly arises due to poor control over undesired amplification close to ambient temperature after initiation.^17^ Similarly, heterogeneity in amplification kinetics due to poor diffusion in the polyethylene glycol (PEG)-rich RPA reaction mix leads to large variability. To bypass these issues and reduce interference from non-specific amplicons, a recent report presented a competitive RPA scheme where a reference DNA competes with the target for primers and amplification enzymes.^18^ Amplicon levels compared to the reference was employed to demonstrate quantification over five log orders. However, such ratiometric techniques require optimized reference templates and additional probes to quantify the reference DNA and hence limiting its ease of implementation.

Development of INAATs can benefit from mathematical models based on mechanistic understanding of the reaction pathway that can identify the parameters controlling the kinetics and yield of target amplicons.^19–21^ For example, a kinetic model for RPA described how mixing is a key limitation controlling the amplification reaction^20^ but the model mostly focused on the initial reaction conditions. A common observation with RPA reactions has been the low and variable yield of amplicons compared to PCR though the underlying factors contributing to this remain uncharacterized.

In this work, we developed a kinetic model for RPA using insights from fluorescence imaging and real-time monitoring of RPA reactions. We show that rapid DNA polymerization progressively leads to formation of DNA amplicon clusters that limits further amplification. A key prediction from our model is that the yield of the RPA amplicons would be dependent on the initial target DNA concentration as the amplification rate becomes comparable to DNA cluster formation rate. We experimentally validate this and report that apart from the threshold time, the amplicon yield can display a linear relationship with log of starting DNA concentration under such conditions. We optimized the RPA reaction to identify the conditions that increase sensitivity and resolution of our estimation based on final RPA amplicon yield as well as demonstrate the robustness of the method. Our semi-quantitative endpoint (qe)RPA can estimate target DNA amount over five log orders with a detection limit of 100 molecules. Importantly, this mitigates the need for real-time monitoring for DNA quantification, lowers background amplification and enables the use of DNA intercalating dye for quantification. Finally, we demonstrate that qeRPA can determine dengue viral load from clinical serum samples over a wide concentration range consistent with corresponding qPCR estimates.

## Materials and Methods

### Nucleic acids for amplification

We use the plasmid pRL02 as template for both standardisation assay and standard curve generation. It codes for partial segments of the Dengue virus serotype 2 (DENV2) genome (NCBI GenBank: AY037116.1, positions from 1-403, 10120-10723) inserted in a pUC57 backbone. pRL02 plasmid was isolated (Qiagen plasmid isolation kit) from DH5*α E. coli* and eluted in nuclease-free water.

For viral load quantification, DENV2 viral RNA was isolated from 250 *μ*l of left-over serum (collected from patients who tested positive with Dengue NS1 antigen test, J. Mitra Co.) in the Dengue Consortium Study in Bangalore, India. Informed consent from patients was received as per study protocols approved by the Institute Ethics Committees where samples were collected, and assays were performed. Extracted RNA (Qiagen viral RNA purification kit) was eluted in nuclease-free water as per manufacturer’s instructions. All samples were serotyped as DENV2 using real-time RT-PCR kit (Geno-Sen’s DENGUE 1-4 Real Time PCR Kit). The viral RNA from patient samples were reverse transcribed with the primer DENV-CRP with the Superscript IV reverse transcriptase according to manufacturer’s instruction (Invitrogen). All template nucleic acids were quantified using Qubit-4 fluorometer when not mentioned otherwise.

### Primers

Primers (FW-535, RW-837) targeting the DENV2 capsid gene (GenBank: AY037116.1, positions from 98-403) were designed and used for RPA standardisation assays. For viral load quantification from clinical samples, separate RPA primers (DENV-CFP4, DENV-CRP) were designed for the conserved regions of the capsid gene based on fifteen India specific DENV2 genomes sequences (NCBI GenBank). Alignment of the sequences was done in Ugene software, and the primers were designed using the IDT oligo analyzer tool. The detection probe was designed according to the manufacturer’s recommendation targeting the capsid region of DENV2 with TAMRA dye and BHQ-2 quencher. Designed primers conform to the following criterias: i) 30-35 bases in length ii) 30-70% GC content iii) Minimum Gibbs free energy (ΔG) allowed for hairpin structures, homodimer and heterodimer formation is −4 kJmol^-1^. All the primer and probe sequences are listed in Table S1.

### qPCR assay

cDNA from reverse transcription of DENV2 clinical viral RNA was quantified using qPCR with the KAPA2G Robust HotStart ReadyMix, 500 nM of primers (DENV-CFP4 and DENV-CRP), 2.04 *μ*M of EvaGreen dye (Biotium), and cDNA template (pRL02). The qPCR reaction (Bio-Rad, CFX96 Touch) was performed as: 95 °C for 3 minutes, followed by 35 cycles at 95 °C for 15 seconds, 66 °C for 15 seconds, and 72 °C for 20 seconds with a final extension at 72 °C for 30 seconds. The real-time trajectories obtained from the qPCR thermocycler (Bio-Rad, CFX96 Touch) is analysed using custom written MATLAB script for quantification by building a standard curve with log(copy number) and threshold cycle of standard template (pRL02).

### RPA reaction

All RPA reactions were carried out using the TwistAmp® Basic kit (TwistDX). For the unmodified RPA reaction, one RPA enzyme pellet was resuspended in 29.5 *μ*l of rehydration buffer and supplemented with 300 nM of primers (FP-535 and RP-837), and 2.04 *μ*M EvaGreen dye. The mixture is then vortexed, and centrifuged, followed by the addition of template and 14 mM magnesium acetate to a total volume of 50 *μ*l. The reaction is kept on ice till start of RPA. The RPA mix was incubated at 39 °C when not mentioned otherwise and was monitored in real-time (Bio-Rad, CFX96 Touch). Optimized qeRPA reaction consists of one RPA pellet resuspended to a final volume of 100 *μ*l with 300 nM primers, supplemented with additional reagents to prepare the qeRPA cocktail comprising of 2.4 mM ATP, 0.24 mM dNTP, 2.04 *μ*M EvaGreen dye, 14 mM Magnesium acetate, template nucleic acid and TD buffer (25 mM Tris pH 7.9 and 3 mM DTT). These modified qeRPA reaction conditions were used when not mentioned otherwise. To compare the effect of detection strategy, EvaGreen DNA dye was replaced with RPA fluorescent probes (DENV2-CAP-probe at 0.12 *μ*M) with 100 U Exonuclease III (NEB), keeping all other reagents same. To test RPA specificity, the pRL02 plasmid in presence of varying concentrations of mammalian genomic DNA (extracted from Vero E6 cells, ATCC CRL-1586) was used as the template.

### Model for RPA dynamics

The major steps in the RPA reaction considered in our model are depicted in Figure 2A. The model simplifies the RPA reaction with insights from experiments and molecular steps are lumped to capture only the key features of the amplification process. Some of the model assumptions are: i) The reactants such as ATP, dNTP, primer, enzyme concentration are not limiting. This assumption is likely not valid at the later stages of the amplification (as discussed later). However, it is reasonable since we observe insignificant increase in amplicons after formation of clusters. ii) DNA proteins bind to the primers quickly. This is justified by usually high association rates observed for recombinases and DNA polymerases^22–24^ and high concentration of enzymes and primers in the reaction mix.

**Figure 1:**
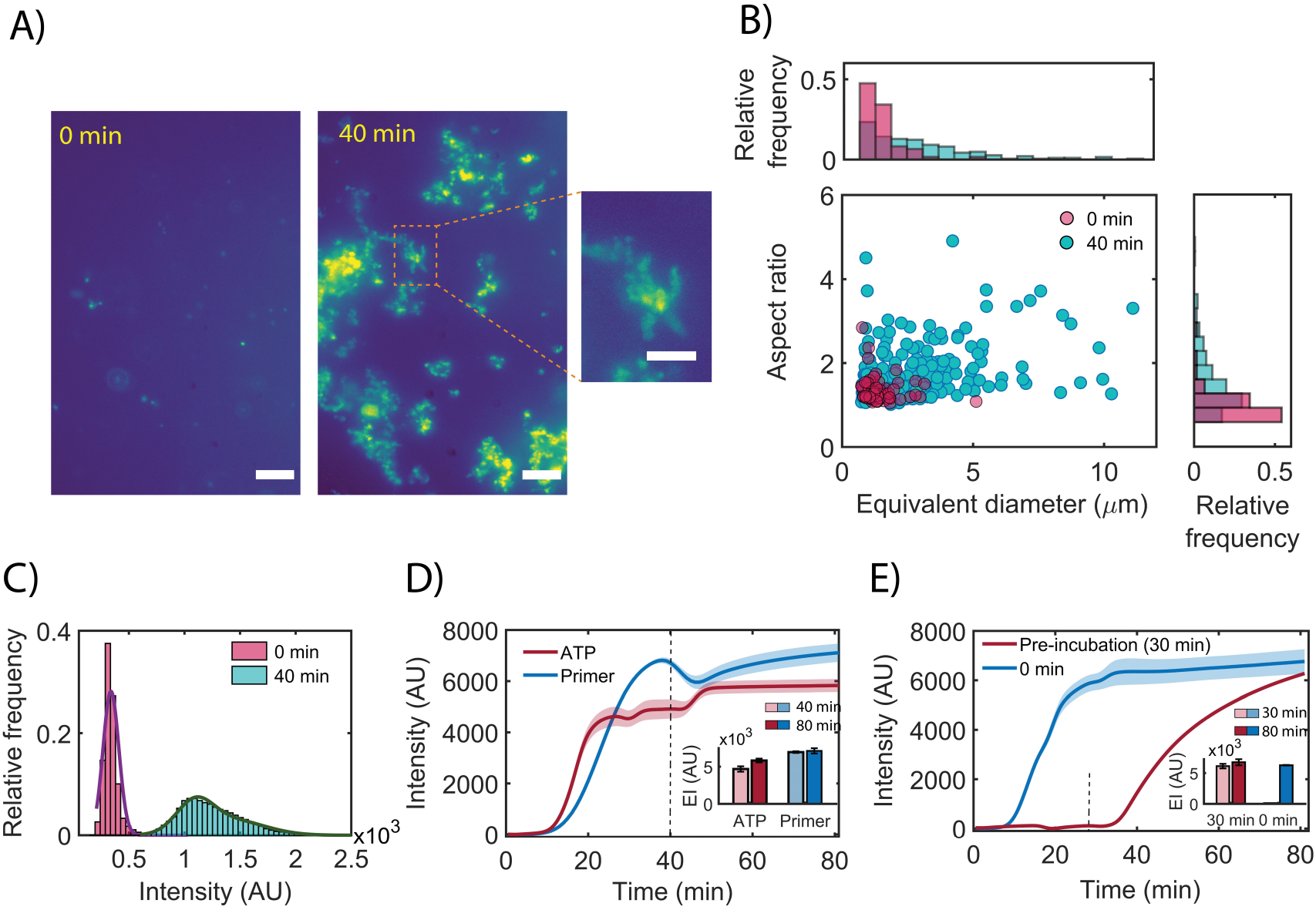
RPA cluster formation (A) Fluorescent images of RPA amplicons before (t=0 minutes) and after 40 minutes of amplification (visualized in presence of EvaGreen dye). Zoomed image highlights the branched and filamentous architecture of the nucleoprotein complexes formed. Scale bar, 10 *μ*m and 5 *μ*m (for the zoomed image). (B) Scatter plot and marginal histograms of the aspect ratio and equivalent diameter of RPA clusters. (C) Pixel intensity histogram of RPA clusters before and after amplification. The solid smooth line represents the Gaussian fit to the intensity histograms. (D) Real-time trajectories of RPA (10^5^ DNA copies) followed after further incubation post addition of fresh ATP and primers at 40 mins (marked as dashed line). Inset bar plot represents the fluorescence intensity at 40 mins and 80 mins from the time traces. (E) Real-time amplification profile for 10^5^ DNA copies when primers and template are added at the start of reaction or after a 30 min preincubation (dashed line). Inset bar plot represents the fluorescence intensity at 30 mins and 80 mins for both cases.

**Figure 2:**
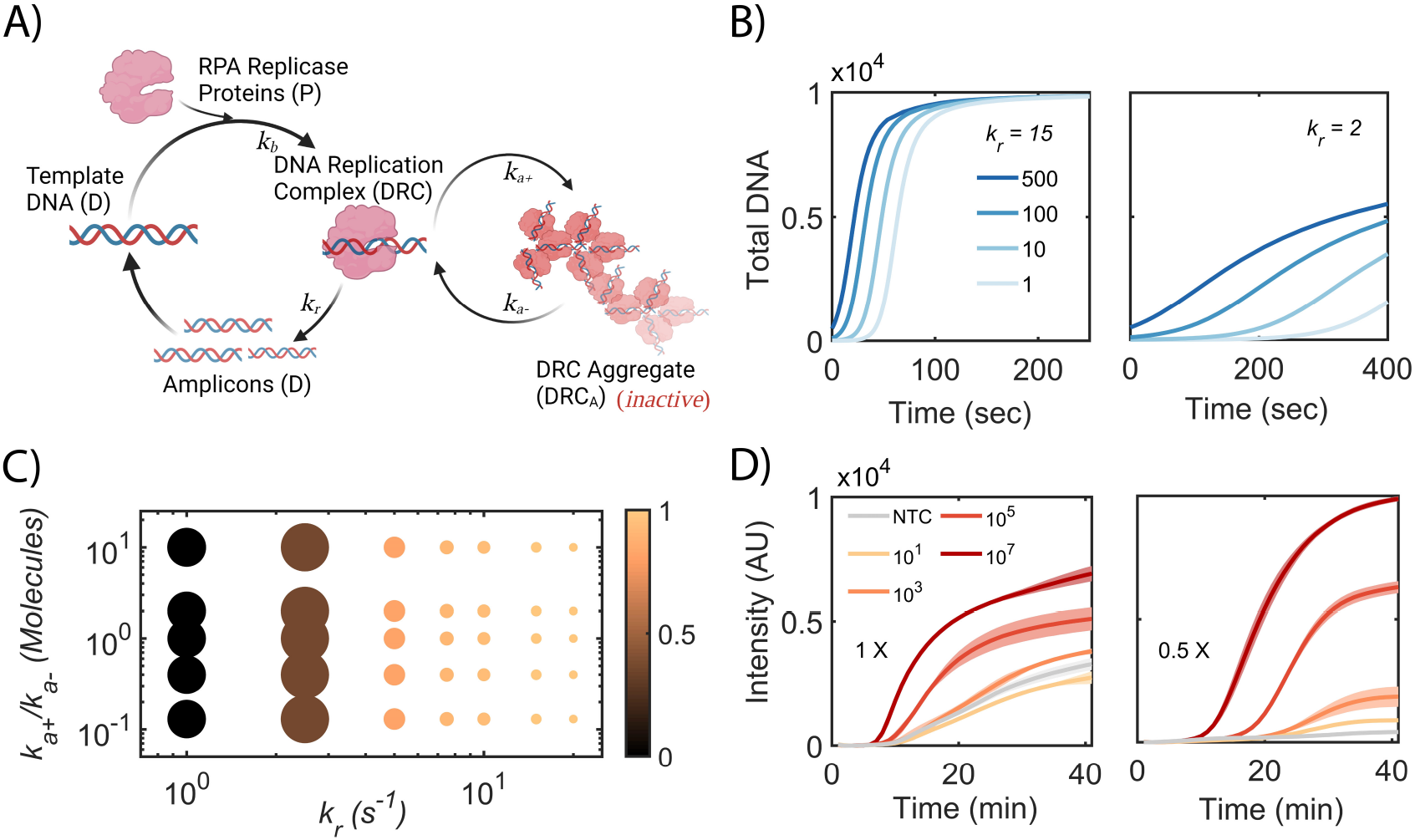
Model for Recombinase Polymerase Amplification (A) Schematic of the RPA model. (B) Model predictions for total DNA (*D_all_*) dynamics for different target DNA concentrations (Left panel for *k_r_* = 15 s^-1^ and right panel for *k_r_* = 2 s^-1^). (C) Variation in amplicon yield and separation between amplified DNA as a function of amplification kinetic rate constants (*k_r_*) and aggregation equilibrium (*k*_*a*+_ / *k*_*a*-_) is shown. Colorbar depicts the magnitude of amplicon yield for 1 copy of target DNA and size of the circle denotes the slope of amplicon yield between 100 and 1 DNA copy. (D) Real-time RPA trajectories for different reaction conditions. Left panel with manufacturer recommended concentration (1X) and right panel with 2-fold dilution of the recommended concentration. The shaded region represents the standard error (SE) from triplicates.

The RPA reaction is represented by three distinct steps. First, the pre-amplification step includes the recombinase-primer complex binding to the target dsDNA (D) and binding of the polymerase to the above complex generates the DNA-replication complex (DRC) (Figure 2A). Second, DRC, upon polymerization, generates new amplicon DNA. Third, DRC complex can reversibly generate larger polymerization-incompetent aggregates called DRC aggregate (DRCA) due to condensation of DNA and proteins during the reaction.

The rate equations used to simulate the kinetics of the reaction are given as follows:

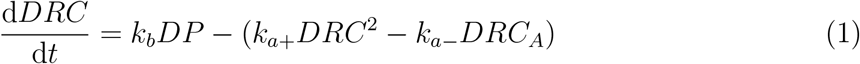

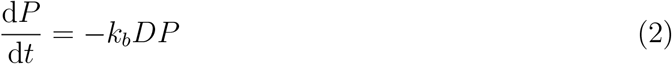

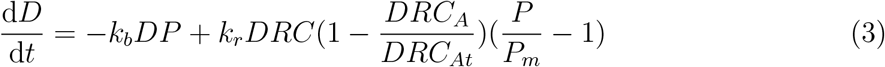

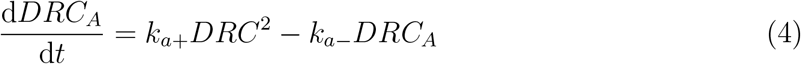

where *D, DRC* and *DRC_A_* denote the number of free dsDNA, the DNA-protein complex for polymerization, inactive form of DNA-enzyme aggregate and *P* represents the DNA proteins in the reaction mixture. *k_b_* and *k_r_* are the rate constants for the DRC formation and polymerization of new DNA, respectively. *k*_*a*+_ and *k*_*a*-_ are the rates for formation and recovery from the inactive *DRC_A_* state. We assume *k_b_* to be high (100 Molecule^-1^s^-1^) consistent with tight binding observed for most DNA binding proteins in RPA. In comparison, *k*_*a*+_ and *k*_*a*-_ is taken as 0.001 Molecule^-1^ s^-1^ and 0.001 s^-1^, respectively. We also impose two constraints to match the reaction conditions. DNA amplification reaches saturation when it either hits the threshold level of *DRCA* (*DRCAt*) or when most of the proteins are used up (implemented as the minimal protein level required for successful polymerization (*P_m_*)). Both control the effective polymerization rate in a logistic fashion (second term in Eq 3). We assume the levels of *DRC_At_* to be high and use rough estimates (20000 molecules) as limited experimental data is available at present. Similarly, the reaction stalls as protein levels drop and approach *P_m_* (set as 10000 molecules). *D_all_* (= *D* + *DRC* + *DRC_A_*) represents the total DNA in the reaction mixture and it is plotted for comparing the amplification kinetics in RPA. Change in these parameters influence the final range of amplicon yields and time to saturation, however, the overall amplicon dependence on initial level of target DNA is reproduced.

### Analysis of RPA amplification trajectories

The real-time amplification kinetics for RPA reactions was extracted and analyzed using the custom-written MATLAB script. Briefly, the raw fluorescence time trajectories obtained from the thermocycler was fit to a four-parameter logistic equation (Eq 5) after background subtraction (background is set as the intensity value at t = 0).

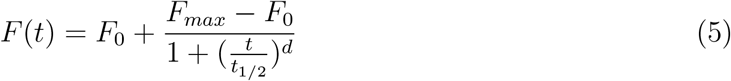

where *F*(*t*) represents the fluorescence intensity as a function of time (*t*). *F_max_* and *F*_0_ represent the endpoint intensity at 40 minutes and baseline intensity, respectively. The variables *t*_1/2_ and *d* denote the point of inflection (time point corresponding to half of the intensity difference between *F_max_* and *F*_0_) and the steepness (or slope) of the curve at point *t*_1/2_, respectively (Figure 3B). From these parameters, threshold time point (TP) is derived as the convergence point of P1 (tangent to *F*_0_) and P2 (tangent at *t*_1/2_) corresponding to the linear part of amplification) as given by Eq 6 which is analytically derived by solving the line equations for P1 and P2 (Supplementary note S2).^25^

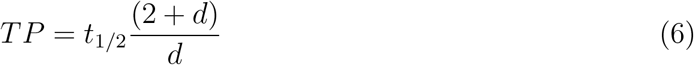

**Figure 3:**
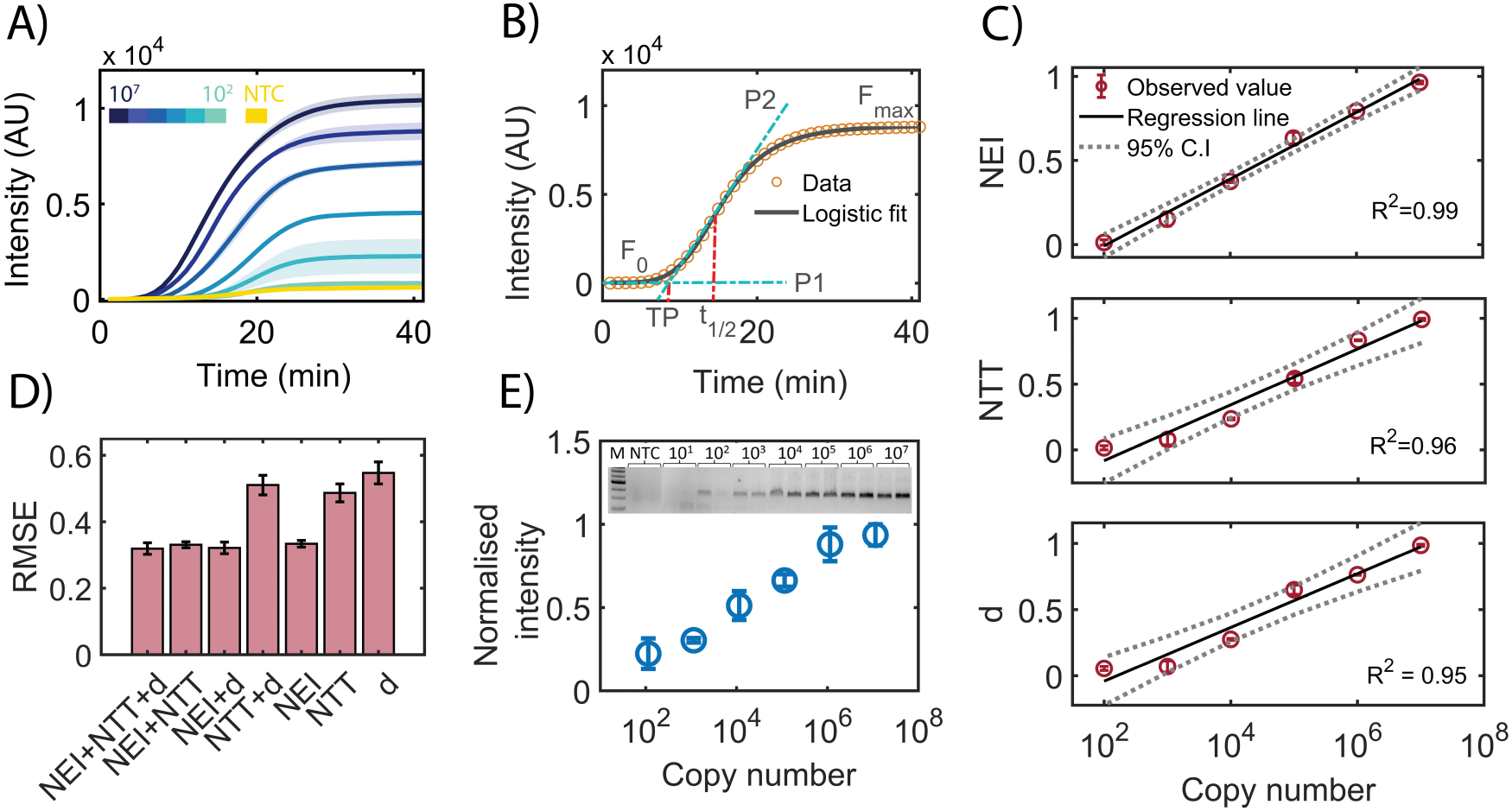
Estimating target nucleic acid concentration using qeRPA (A) Real-time RPA trajectories for serially diluted target DNA ranging from 10^7^ to 10^2^ molecules are plotted. The shaded region represents the SE from duplicate experiments. (B) RPA real-time trajectory is fit to a four-parameter logistic function (Eq 5). (C) Linear regression analysis of NEI and kinetic parameters is shown. The error bar represents the SE from duplicate experiments and 95% CI is depicted between the dashed curves (D) RMSE from the model fit with mixed (linear and multiple linear) regression models for estimating the relationship to target DNA copy number for different factors is shown. (E) Normalised fluorescence band intensity of RPA amplicons analysed from the gel electrophoresis is shown for different target DNA inputs. Inset represents the gel electrophoresis image.

Finally, we define normalized endpoint intensity (NEI), normalised threshold time (NTT) as shown below.

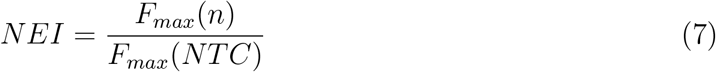

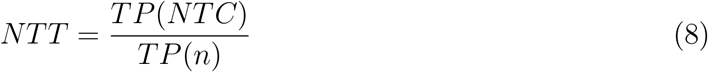

where, *n* and *NTC* denotes the target DNA levels and negative control with no template DNA.

### Imaging and Data analysis

RPA amplified mixture was imaged on an inverted epi-fluorescence microscope to capture the fluorescence intensity from EvaGreen dye-bound dsDNA. The imaging glass coverslip surface was treated with dichloro dimethyl silane^26^ (DDS) (Sigma-Aldrich:440272) and further passivated with 2 mg/ml Bovine serum albumin (BSA, NEB:B9000S) to prevent non-specific binding of proteins and DNA to the surface. The sample was excited with a 488 nm laser and the emitted fluorescence was captured from different axial plane on a sCMOS camera (ORCA FLASH 4.0) with an 100X (Olympus, NA:1.45) objective. The excitation and emission paths included a longpass dichroic filter (511 nm), and a bandpass filter 538/40 nm (Semrock), respectively.

The captured fluorescent images are analysed using a custom written MATLAB script. Initially, the algorithm finds the maximum intensity projection from different z-stacked images of the RPA mix. Background intensities are calculated using a dilated image and subtracted from the original image. Intensity threshold is calculated from the fitting of pixel intensity histogram to a two component Gaussian mixture model. A binarized image is generated with the thresholding applied based on this cutoff and the connected components are merged to identify the discrete bright features (DNA clusters) in the image. Equivalent diameter (=4 * *Area/Perimeter*) and aspect ratio (= *Major axis length/Minor axis length*) is estimated for the detected DNA clusters from the binary image.

## Results and discussion

### Nucleoprotein cluster formation during RPA

RPA proceeds at 39±2 °C in the presence of high concentrations of co-solutes that include various DNA binding proteins and PEG polymer. The commercial RPA cocktail is optimized for rapid DNA amplification from low target DNA levels. However, under the high concentrations of proteins, PEG, and DNA amplicons, the reaction conditions can lead to condensation and spontaneous formation of nucleoprotein clusters.^27^ We speculated that such aggregation would be driven by the local accumulation of new amplicons fueled by rapid DNA amplification during RPA. Indeed, formation of RPA clusters that correlate with starting DNA concentration has been reported recently.^28^

To examine the formation of RPA clusters, we visualize the DNA products using an intercalating dye (EvaGreen) during RPA under high magnification (100X magnification). Upon amplification, we observe emergence of irregular aggregates with a broad size distribution and spanning up to 10 *μ*m in effective diameter (Figure 1A, B). This is in contrast to the limited number of uniform and small puncta observed for target plasmid DNA before amplification. There is a 4-5 fold higher pixel intensity in the clusters compared to plasmid DNA suggesting higher density of DNA in the aggregates (Figure 1C). Since recombinase and single stranded protein (SSB) involved in RPA are known to interact and form filaments on DNA,^29^ we attribute these DNA-rich aggregates to nucleoprotein clusters formed in conjunction with the recombinase and SSB complexes on DNA.

We speculated that the formation of these nucleoprotein clusters could lead to inhibition of further DNA amplification. Indeed, when we supplement the reaction with additional ATP or primers after formation of the clusters, there is negligible further amplification (Figure 1D). Alternately, to test whether RPA reaction stalls due to loss of protein activity over time, we added primers and template to the RPA reaction after pre-incubating the proteins (at 39 °C for 30 mins). However, in this case, we observe no reduction in final amplicon yield suggesting that loss of enzymatic activity is not limiting further amplification (Figure 1E). This suggests that saturation in RPA reaction is tied to the observed nucleoprotein cluster formation.

### Model for Recombinase Polymerase Amplification

Taking cue from the experimental evidence that suggests limited amplification beyond the formation of clusters, we conjectured that varying the balance between amplification and nucleoprotein cluster formation kinetics can modulate the amplicon yields. To generate a mechanistic insight into the possible outcomes of RPA reaction, we developed a simple kinetic model describing the production of DNA amplicons. We consolidated the RPA reaction into three distinct and broad steps as discussed in the methods section (Figure 2A). Our RPA reaction scheme is captured by the set of four equations (Equation 1-4). We lumped the kinetics of DNA, protein and primer binding in a single rapid molecular binding step with the rate constant, *k_b_.* Similarly, extension and final product release is represented with a growth rate constant, *k_r_*. We also include a by-product DRCA formed from amplicons (rate constant, *k*_*a*+_) as observed experimentally for RPA products (Figure 1A). Here, we consider *k_r_* to be larger than *k*_*a*+_ since clusters were observed only in the latter half of the RPA reaction. We also conjecture that the DRCA complex is inactive but can revert to the active form, DRC albeit at a slow rate represented by rate constant, *k*_*a*-_.

Our RPA kinetic scheme displays a sigmoidal growth dynamics (as observed for the amplicon generation in RPA experiments) when the rate of amplification (*k_r_*) is comparable or larger than aggregation rate (*k*_*a*+_). Additionally, the time to cross a threshold signal level (*τ_d_*) increased logarithmically with the initial DNA levels. This is consistent with previous reports and has been used as a parameter for quantitative estimation of DNA with RPA.^14,30^ However, with a large *k_r_* (≥ 10 s^-1^), we observe identical final amplicon levels irrespective of the initial DNA concentrations.

From our model, we recognized that reduction in the polymerization rate compared to RPA cluster formation can introduce reaction limitations that will manifest in early termination of amplification in RPA. As *k_r_* is reduced, we observe a slower kinetics as well as reduced amplicon yield due to premature inactivation of RPA by aggregation (Figure 2B,C). Once the available proteins drop or DRCA reaches a threshold, RPA stalls and reaches a plateau phase. However, this resulted in an improved separation between the threshold time as well as amplicon levels for different amounts of initial DNA. Evaluating this relationship over a large range for *k_r_* and aggregation equilibrium (*k*_*a*+_/*k*_*a*-_) shows that at intermediate values of k_r_, one can achieve enhanced separation between the amplicon levels without compromising the final yield significantly (Figure 2C). Therefore, apart from the threshold time of amplification, final amplicon yield can be employed for quantitative estimation of nucleic acids in certain RPA reaction conditions.

### Modulating amplicon yield in RPA

To alter the kinetics of the amplification reaction, we conducted the RPA under different dilutions of the reaction cocktail. Further, to reduce costs and avoid slower kinetics associated with RPA probes, we employed a DNA-intercalating dye (EvaGreen) to monitor the RPA kinetics in real-time. We designed RPA primers against dengue virus capsid gene and amplified DNA segments from a plasmid with part of dengue serotype 2 virus genome. When we perform amplification with reduced amounts of RPA enzymes, the kinetics and yield of the RPA is attenuated progressively with reduction till there is poor (or no) yield (Figure S2). However, under partially reduced kinetics (at 2-fold dilution of the RPA mix (0.5X)), the yield of the amplified DNA was monotonically dependent on the initial target DNA concentration as predicted by our model (Figure 2D). This is in contrast to the poor dependence of amplicon yield on initial DNA with the manufacturer’s recommended RPA reaction conditions (termed 1X). Comparison with our model suggests that the commercial RPA kit is optimized to operate in high *k_r_* regime to achieve rapid amplification.

Another advantage of reducing amplification kinetics is the dramatic reduction in the formation of non-specific and primer-dimer products (Figure 2D and Figure S3A) enabling a simpler interpretation of RPA kinetics and amplicon yield data. We also observe heterogeneous amplification with large variation in both threshold time and endpoint yield even for RPA reactions at 2-fold dilution, that is a challenge for quantitative RPA (Figure S3C). However, the variability was significantly reduced when the reaction mixture was mixed thoroughly by pipetting (Figure S3C). Separately, we also verified that the specificity for the template is not compromised in the presence of mammalian genomic DNA under the enforced slower kinetics of amplification (Figure S3B).

We also note that the RPA reactions terminate in experiments beyond 40 mins. This is unlike our model predictions. Even with reduced kinetic rate of amplification, the final amplicon yields would achieve similar levels for all input DNA upon extended incubation. We speculate this discrepancy with our model could arise from near complete hydrolysis of ATP or a still unknown phenomena. In spite of this, data from Figure 1 suggests that cluster formation is the leading cause of stalling further DNA amplification. We also note a moderate increase in the amplicon yields (comparison to recommended enzyme concentration) in some cases (eg. for 10^7^ and 10^5^ copies of DNA in Figure 2D) even when the kinetics is reduced (as is evident from larger threshold times). This we attribute to possible change in the rate of RPA cluster formation effected by the dilution of the PEG and other co-solutes. Nevertheless, this demonstrates that amplicon yield can be monotonic function of initial DNA level in certain RPA reaction conditions.

### Establishing nucleic acid quantification using amplicon yield in RPA

We envisioned that correlative end product formation can be utilised to develop a quantitative RPA without the need for real-time monitoring. For this, we evaluated the real-time RPA amplification trajectories from serial dilution of dengue plasmid (Figure 3A). Using maximum fluorescence intensity (*F_max_* at 40 mins), baseline intensity (*F*_0_), time to reach half of the fluorescence dynamic range (*t*_1/2_), and slope of the amplification phase (*d*) recovered from a logistic fit, we evaluate parameters that can report on initial DNA concentration (Figure 3B).

We estimate normalized endpoint intensity (NEI) that reports on the RPA amplicon yield while normalised threshold time (NTT), and slope of the amplification phase (*d*) that report on the kinetics and efficiency of the amplification. As is evident from the real-time RPA amplification trajectories, normalized threshold time, NTT is linearly correlated to log of starting copy numbers consistent with reports with threshold time^14^ (Figure 3C). Additionally, a positive linear correlation is observed for endpoint amplicon yield (reported as NEI) for a wide range of initial DNA starting from 100 to 10^7^ molecules (Figure 3C). Rate of the amplification phase (captured as *d*) also displays a correlated increase with initial DNA concentration. We derive empirical linear relationships between input DNA and all these factors individually or in combination (Figure 3C). We find that a linear regression model either with a combination of parameters with NEI or NEI alone has the least root mean square error (RMSE) (Figure 3D).

We further validated that NEI was an accurate estimate of target amplicon yields using gel electrophoresis (Figure 3E). Indeed, the amplicon band intensity also replicate the observed linear relationship to as low as 100 copies. Non-specific amplification starts to interfere with target amplification below 100 copies which sets the limit on our ability to quantify accurately below this regime. Neverthless, this shows that amplicon yield is a robust parameter to estimate target nucleic acid concentration over a large dynamic range and it can alleviate the need for continuous monitoring of RPA. We name this approach, quantitative endpoint RPA (qeRPA). We observe the same increasing trend visually for amplicon yield (EvaGreen fluorescence intensity) in the reaction vial post amplification (Figure S4). Therefore, qeRPA is also amenable to visual qualitative inspection of amplicons despite the overall reduction in yield and slower kinetics.

We also verify that the RPA kinetics and the endpoint amplicon yield for qeRPA can be captured with specific DNA based RPA probes. Consistent with DNA dye based qeRPA, we observe that final amplicon yield is a function of initial target DNA level (Figure S5). However, a slower kinetics was observed for probe based qeRPA suggesting that probe binding and processing is the rate limiting step. On the other hand, increased specificity of probes results in better signal to noise ratio for amplicon yield (inset in Figure S5). Therefore, endpoint amplicon based qeRPA can be applied irrespective of choice of detection strategy for amplicon yield. Combined with challenges in designing efficient RPA probes and associated costs, we argue that DNA dye based detection is a better reporter for RPA dynamics and hence we chose EvaGreen for further characterizing qeRPA.

### Robustness of qeRPA under various reaction conditions

Since the modifications to the RPA reaction enabling qeRPA can compromise the reaction under diverse conditions, we examined whether and when qeRPA fails. To determine the robustness of qeRPA, we examined the effect of various factors (temperature, primer and magnesium levels) on the end product yields and subsequent quantification. For a quick evaluation of qeRPA under various conditions, we compare dengue amplicon yields (reported for 1000 DNA copies) and the extent of separation in amplicon yield between low and high DNA concentration regimes (estimated as the slope between amplicon yields at 1000 and 1 million copies of target DNA). These two criteria define the detectable signal (sensitivity) and the resolution for DNA estimation by qeRPA and are depicted on two orthogonal axis for each condition (Figure 4). The recommended operating temperature ranges from 37 - 42 °C for RPA. At temperatures above 40 °C, either the amplicon yield and separation between copy numbers is compromised but all conditions (apart from 42 °C which displays poor amplicon yield) should perform comparable to each other (Figure 4A). At all the temperatures below 42 °C, qeRPA generated specific products and fewer non-specific products as observed by gel electrophoresis (data not shown).

**Figure 4:**
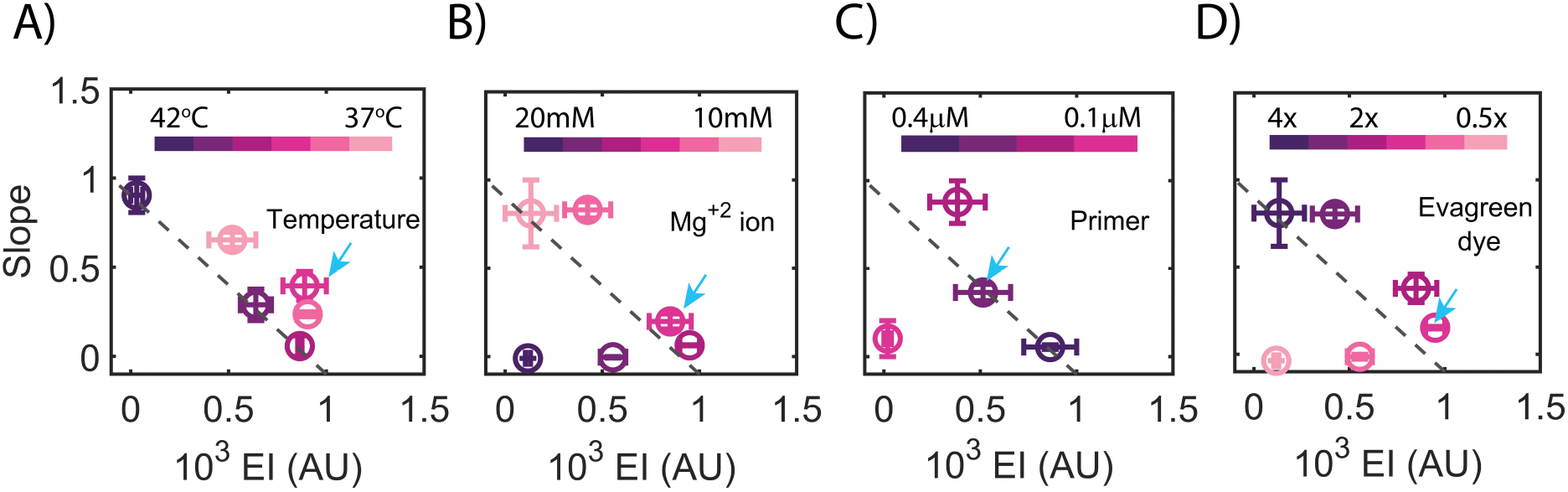
Efficiency of qeRPA reaction under different conditions. The x-axis and y-axis represents the normalised endpoint intensity of 1000 DNA copies and slope of NEI vs log(copy number) for each dataset. qeRPA performance under different A) temperature: 42 °C to 37 °C with an interval of 1 °C B) Mg^+2^ ion: 20 mM, 18 mM, 16 mM, 14mM, 12mM, and 10 mM C) primer concentration from 0.4 *μ*M to 0.1 *μ*M with 0.1 *μ*M interval D) EvaGreen dye: 4x, 3x, 2x, 1.5x, 1x and 0.5x. 1x corresponds to 1.36 *μ*M. The blue arrow in each plot point to the concentration or temperature used in the optimized qeRPA. Dashed grey line is a visual aid to demarcate qeRPA performance with conditions above the line considered acceptable. Error bar represents the SE from duplicate experiments.

Magnesium serves as a co-factor for UvsX recombinase and Sau DNA polymerase in RPA and affects the stability of duplex DNA. Between 12 to 16 mM, RPA generates specific amplicons, detectable amplicon yields, and optimal separation for quantification with qeRPA. However, at 10 mM Mg^+2^, RPA amplicon yield is poor and at concentrations above 16 mM, there is poor resolution in distinguishing DNA levels(Figure 4B). Therefore, the optimal window of magnesium concentration is consistent with the recommended conditions for conventional RPA.^31^ The optimum concentration of primer is critical for specific product formation compared to template-free primer extension products. Moreover, recombinase-bound primer levels are expected to define the amplification rate in RPA and hence recombinase-primer stoichiometry is a key parameter. Figure 4C suggests 0.2 *μ*M primer concentration as the best performing condition with a high value for slope and endpoint intensity. However, with increasing primer concentration (from 0.2 *μ*M to 0.4 *μ*M), the endpoint intensity increases but with reduced separation between the amplicon levels. This is expected from increase in the amplification rate (*k_r_*) consistent with our model (Figure 2B). At lower concentrations (0.1 *μ*M), the amplicon yield is less predictive for quantification due to poor amplification. We also examined the effect of the DNA dye on qeRPA resolution and yield. EvaGreen dye at 2.04 *μ*M - 4.08 *μ*M (corresponding to 1.5X - 3X of recommended for qPCR) for real-time detection in qeRPA displayed high signal intensity and resolution. Lower dye concentrations (≤1X) do not provide sufficient distinction between the different DNA levels due to poor dye:DNA stoichiometry. On the other hand, higher dye concentration (4X) results in reduced amplicon yield indicating the inhibitory effects of EvaGreen dye on RPA (Figure 4D).

Due to the limitations placed on the amplification kinetics, we identify an overall inverse relationship between amplicon yield and resolution of qeRPA (Figure 4). Nevertheless, we argue that qeRPA performance is generally robust over conditions consistent with RPA protocols. Based on this analysis, we recommend incubation at 39°C for 40 mins with 2-fold dilution of RPA mastermix, 14 mM magnesium, 0.3 *μ*M primers and 2.04 *μ*M EvaGreen dye for qeRPA analysis.

### Quantification of Dengue viral RNA from patient serum

To demonstrate the clinical applicability of qeRPA, we next determined the dengue viral load in patient serum samples with active dengue infection (Figure 5B). With a new set of primers designed against the conserved regions in the capsid gene of the dengue serotype 2 genomes prevalent in India, we confirmed specificity and sensitivity for qeRPA (Figure S6).^32^ We re-evaluated qeRPA with these primer sets and determined optimal conditions to be 0.3 *μ*M primers, 14 mM Mg^+2^, and 40 °C which was similar to our previous conditions confirming previously characterized conditions for qeRPA.

**Figure 5:**
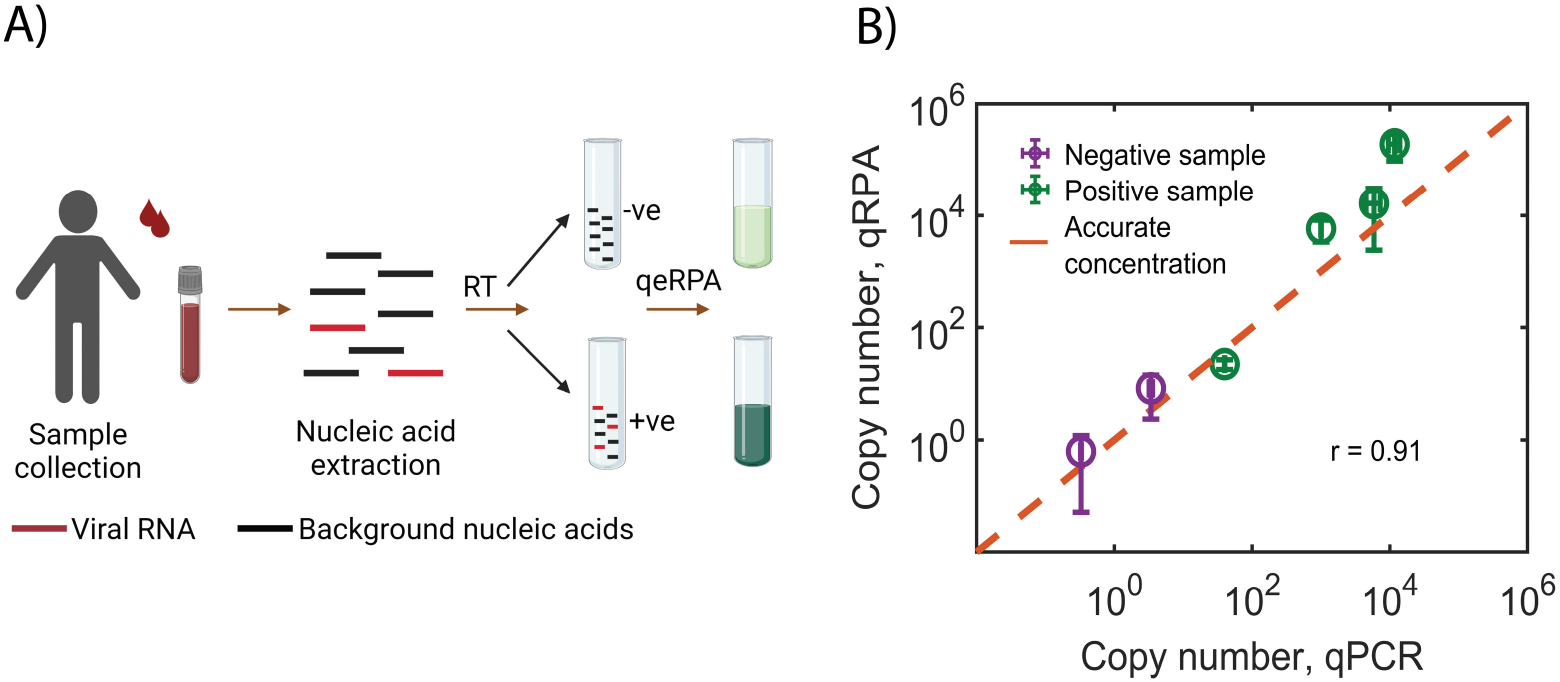
Viral RNA quantification from clinical samples. (A) Blood from dengue infected patients are collected and RNA extracted is reverse transcribed (RT) to cDNA. The samples with dengue viral cDNA shows an increased fluorescence signal compared to the negative samples devoid of viral cDNA. (B) Comparison of the copy numbers calculated by two methods qPCR and qeRPA. Error bar represents the SE from triplicate experiments

Viral cDNA generated from purified serum RNA are quantified by qPCR and compared to qeRPA estimates. *C_t_* values for qPCR with the RPA primers for two control samples (C1 and C2) were >31 and for all the four positive samples (P1-P4) tested were <29. We estimated the viral cDNA copy numbers from qPCR using the standard curve generated with pRL02 plasmid (Figure S6A). Similarly for qeRPA, the standard curve is generated using NEI after amplification (Figure S6B). Viral nucleic acids levels quantified with qeRPA are concordant with the qPCR estimates for all the positive samples (Figure 5B). The qeRPA was able to predict the copy numbers reliably (r = 0.91) and the largest variation observed at the highest viral load was within a log order of magnitude from the qPCR estimate. Interestingly, we could detect specific amplification of the dengue amplicon with qeRPA even below 100 copies (sample-P4,Figure S6C). Even for control samples (with no specific target band), the level of non-specific amplification was comparable between qPCR and qeRPA. This suggests that further RPA primer optimization can improve the detection limit in qeRPA. Nevertheless, the qeRPA method can accurately estimate the concentration of viral nucleic acids from clinical samples.

## Conclusion

In this work, we developed an optimized quantitative isothermal method that bypasses the requirement for continuous monitoring. The method was able to accurately detect and quantify target nucleic acids within ~40 minutes of reaction over a large dynamic range and is robust over most reaction conditions. This method is less susceptible to errors in estimates of threshold time based quantification as the background RPA reaction can proceed at room temperature.

The qeRPA approach presented here has a dynamic range of 10^5^ with a demonstrated detection limit of 100 molecules. Further improvisation in the primer design or RPA reaction mix composition can reduce non-specific products, increase amplicon yields and their separation in the qeRPA assay. Alternatively, usage of target-specific probes can also increase the specificity and sensitivity of qRPA assay.

## Acknowledgement

The authors thank Twinkle Gupta and Suraj Jagtap for their help with the primer design for Indian dengue genomes and Saranya Marimuthu for sharing the plasmid pRL02. This work was supported by Department of Science and Technology, DST-SERB grant CRG/2018/004266 (RR), Department of Biotechnology, DBT Biodesign-Bioengineering Initiative BT/PR13926/MED31/97/2010 (RR) and PhD fellowship from Indian Institute of Science (PV).

## Supporting Information Available

Supplementary information is given in supplementary information document.

