## Supplementary file for "Nucleic acid quantification with amplicon yield in recombinase polymerase amplification"

### ELECTRONIC SUPPLEMENTARY INFORMATION (ESI) for Nucleic acid quantification with amplicon yield in Recombinase Polymerase Amplification

Priyanka Valloly[a], and Rahul Roy*[a,b]

**a** Department of Chemical Engineering, Indian Institute of Science, Bangalore, Karnataka, India - 560012

**b** Center for BioSystems Science and Engineering, Indian Institute of Science, Bangalore, Karnataka, India - 560012

#### Content

Fig S1: RPA amplicon yield and sensitivity as function of binding and aggregation kinetics

Fig S2: Modulating RPA polymerization kinetics

Fig S3: Sensitivity, specificity and effect of mixing in qeRPA

Fig S4: Final amplicon yield quantification in reaction tubes with an imager

Fig S5: RPA kinetics using probes and EvaGreen dye

Fig S6: Standard curve for viral RNA quantification from clinical samples

Table S1: List of primers and probes

Table S2: Parameters used in the RPA model simulation

Note S1: Accession numbers of the DENV2 genome sequences from NCBI GenBank database

Note S2: Analytical solution for threshold time point (TP)

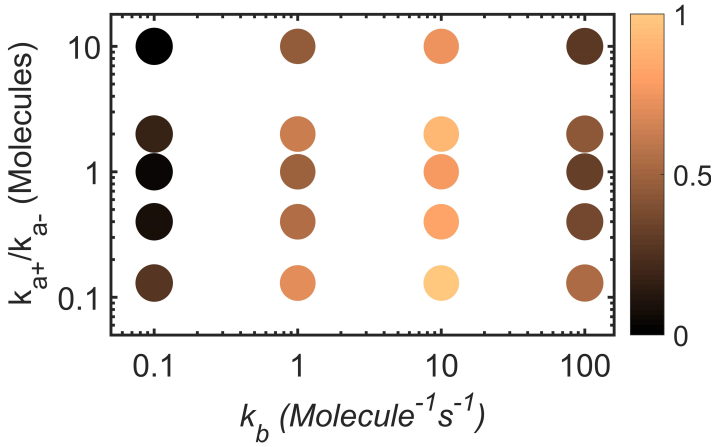


**Fig S1. RPA amplicon yield and sensitivity as function of binding and aggregation kinetics**

Size of the circle signifies the slope of amplicon yield between 1 and 100 DNA copies. Color bar indicates the normalised amplicon yield for 1 copy of DNA template. Unlike observed for $k_r$, changes in $k_b$ do not modulate the level of separation between amplicon yields while the amplicon yield drops significantly at high as well as low extreme values for $k_b$.

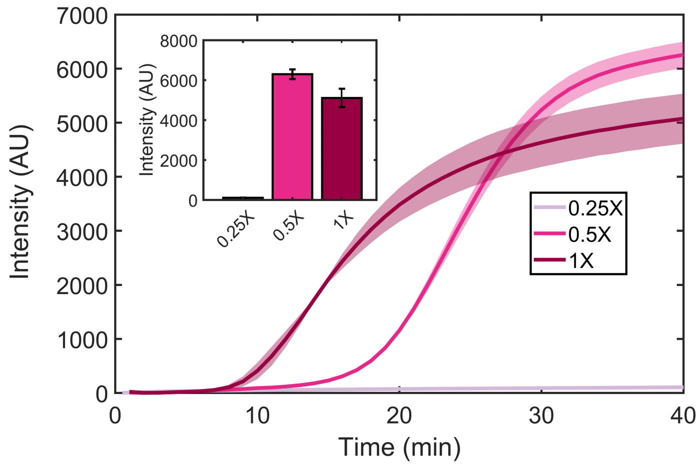


**Fig S2. Modulating RPA polymerization kinetics**

Real-time trajectories of RPA with different dilutions of RPA cocktail with $10^5$ copies of pRL02 plasmid is shown. At 2-fold dilution, the kinetics is slower but marginally higher yield of amplicons is achieved. Further dilution (4-fold), reduces the kinetics dramatically such that no amplification is observed. Inset bar plot represents the endpoint intensity for different enzyme concentrations. The shaded region and error bar (inset graph) represents SE from duplicate experiments.

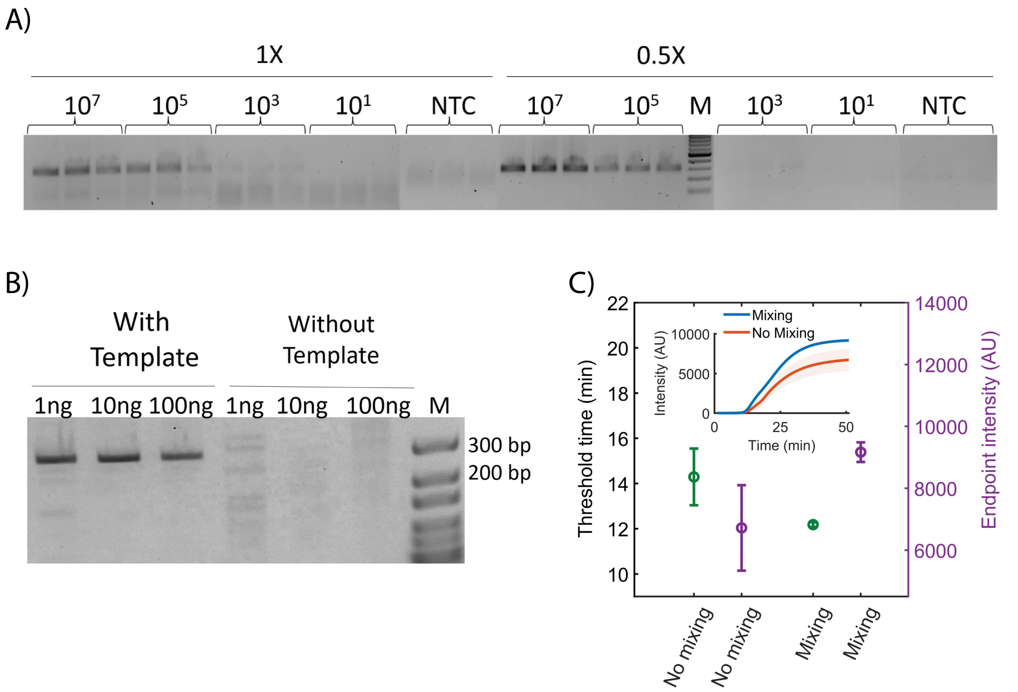


**Fig S3. Sensitivity, specificity and effect of mixing in qeRPA.**

(A) Comparison of RPA reaction yield and non-specific product formation with 1X and 0.5X enzyme concentration for different template copy numbers on 2% agarose gel stained with ethidium bromide. 'NTC', and 'M' represent negative control, and marker respectively. (B) Specificity of designed primers in presence of a different concentration of mammalian genomic DNA along with the $10^5$ copies of target DNA. (C) Efficiency of RPA reaction on mixing. The threshold time and endpoint intensity observed in RPA with and without adequate mixing are shown with SE from duplicate experiments. Inset curve represents the real-time trajectories for RPA performed with inadequate mixing (red curve) and with mixing (blue curve). Inadequate mixing leads to large variations in threshold time as well as amplicon yield.

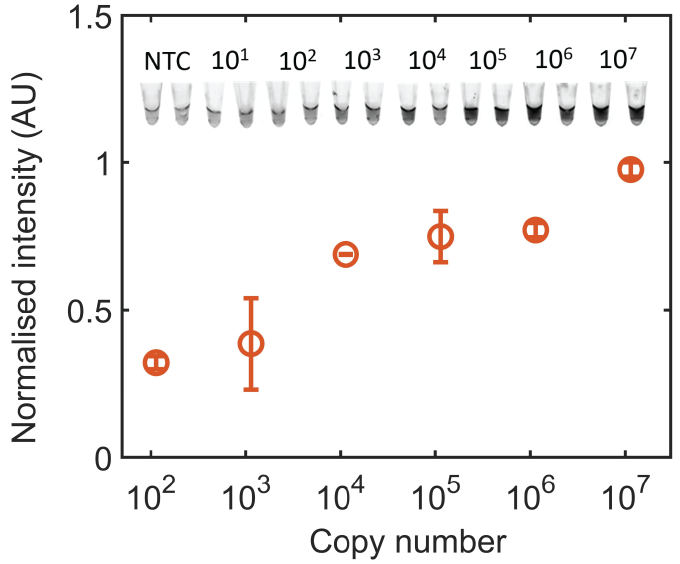


**Fig S4. Final amplicon yield quantification in reaction tubes with an imager**

Intensity quantified from EvaGreen dye bound amplified products from the tube images (as shown in inset) after RPA reaction are reported. The inset of the graph represents the tube images of amplified product taken with a fluorescent imager (at 365nm excitation and 400-450 nm emission) at different starting DNA copy numbers with NTC as the no-template control.

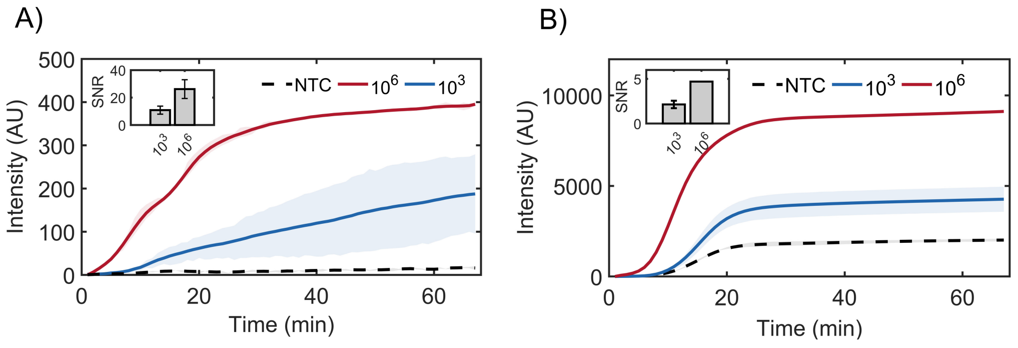


**Fig S5. RPA kinetics using probes and EvaGreen dye**

Real-time fluorescence trajectories captured with A) Dengue-specific probes and B) EvaGreen dye.The inset depicts the ratio of fluorescence signal of template DNA to background (NTC) for each concentration. The shaded region and error bar represents the SE from duplicate experiments.

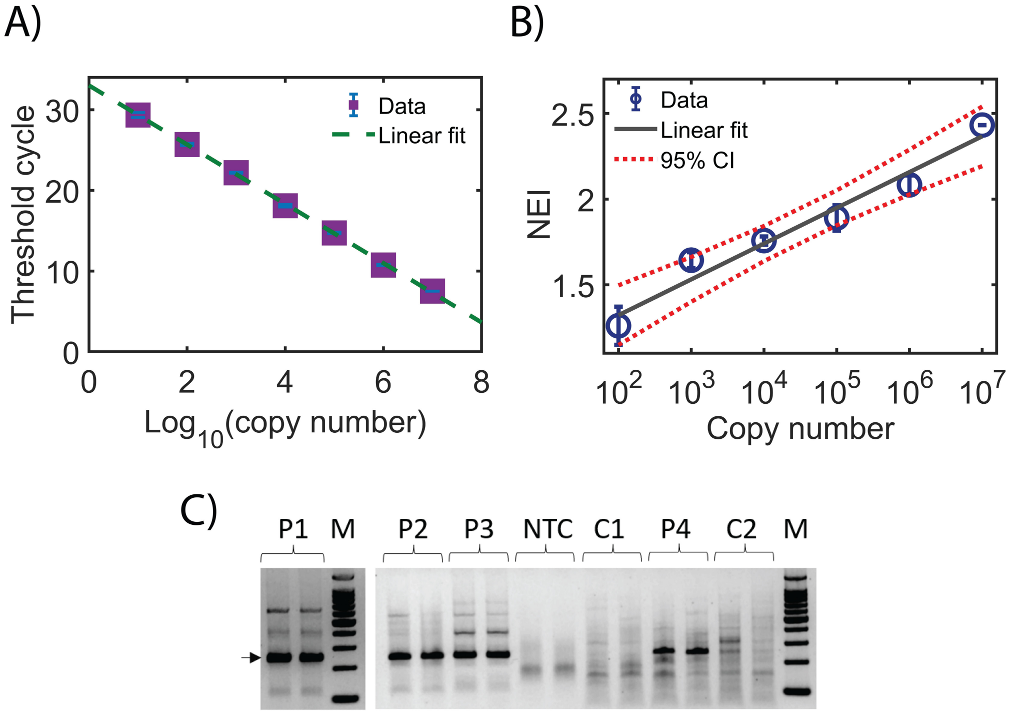


**Fig S6. Standard curves for viral RNA quantification from clinical samples.**

(A) Real-time qPCR standard curve with EvaGreen dye with pRL02 plasmid. The error represents the SE from duplicate experiments. (B) Standard curve for qeRPA with copy number on the X-axis and NEI on the Y-axis. The error bar represents the SE from triplicates. (C) Gel elctrophoresis analysis of RPA of clinical samples. P1-P4 represent the positive samples for dengue infection and C1-C2 represents the control sample without dengue infection. 'NTC', and 'M' represent negative control, and marker respectively. The arrow indicates the desired amplicon size for RPA with dengue viral cDNA.

| Oligo name | Sequence ($5^{'}$ - $3^{'}$) |
|---|---|
| FW-535 | TGAATAACCAACGGAAAAAGGCGAGAAAAACC |
| RW-837 | CAGTTCTGCGTCTCCTGTTCAAGATGTTCAGC |
| DENV-CFP4 | ACCTTTCAATATGCTGAAACGCGAGAGAAACC |
| DENV-CRP | GCATCCTTCCAATCTCTTTCCTGAACCCTCTC |
| DENV2-CAP-probe | ATTAAAAATTAATTCTCCAGCAGCATTA TCTTACCCTCTC/iFluorT/GC/idSp/AC/iBHQ-idT/TTTGGCGACTCA/3SpC3/ |

**Table S1.** List of primers and probe

| Parameters | Value |
|---|---|
| $k_b$ | 0.1−100 Molecule$^{-1}$s$^{-1}$ |
| $k_r$ | 1−20 s$^{-1}$ |
| $k_{a+}$ | 0.001 Molecule$^{-1}$ s$^{-1}$ |
| $k_{a-}$ | 0.0075−0.0001 s$^{-1}$ |
| $DRC_{At}$ | 20000 Molecules |
| $P_m$ | 10000 Molecules |

**Table S2.** Parameters used in the RPA model simulation

#### S1 Accession numbers of the Indian DENV2 genome sequences from NCBI GenBank database

MG560144, JX475906, JQ955624, JQ955623, MG560143, KU509271, KJ918750, KY427084, JQ922553, JQ922552, JQ922551, JQ922550, FJ898454, DQ448231, JQ922549, KY427085.

#### S2 Analytical solution for threshold time point (TP)

Equation of line $P_1$

$$F_1 = \beta_1 t + \alpha_1 \tag{1}$$

$$\beta_1 = dF_1/dt_{(t=t_{1/2})} = 0 \tag{2}$$

$$\alpha_1 = F_{1(t=0)} = F_0 \tag{3}$$

Equation of line $P_2$

$$F_2 = \beta_2 t + \alpha_2 \tag{4}$$

$$\beta_2 = dF_2/dt_{(t=t_{1/2})} \tag{5}$$

$$\alpha_2 = F_{2(t=m)} - \beta_2 t_{1/2} \tag{6}$$

$$\beta_2 = (F_0 - F_{max})d/(4t_{1/2}) \tag{7}$$

$$\alpha_2 = (F_0(2-d) + F_{max}(2+d))/4 \tag{8}$$

Where $F$,$\beta$,$\alpha$ denotes the intensity, slope and intercept of the corresponding tangent line (P1 and P2). $F_0$, $F_{max}$, $t_{1/2}$ and $d$ indicates the baseline intensity, endpoint intensity at 40 minutes, time to reach half of the fluorescence dynamic range ($F_{max}$ and $F_0$) and slope of the amplification phase respectively. Since both the lines $P_1$ and $P_2$ intersect at TP,

$$\beta_1 TP + \alpha_1 = \beta_2 TP + \alpha_2 \tag{9}$$

Solving for TP,

$$TP = t_{1/2}(2+d)/d \tag{10}$$
